# Evaluating Seqstant LiveGene Analysis in Real-Time Assessment of Metagenomic Next-Generation Sequencing (mNGS) Data from Respiratory Samples

**DOI:** 10.1101/2025.01.24.634829

**Authors:** Sébastien Boutin, Sabrina Klein, Gerold Untergasser, Tobias P. Loka, Suzan Jakob, Elham Khatamzas, Georg Wrettos, Henri Knobloch, Dennis Nurjadi

## Abstract

**Background:** The detection of pathogens causing infections by conventional diagnostic methods can be challenging and next-generation sequencing (NGS) technology offers a promising alternative method. In this study, we evaluated the performance of real-time metagenomic next-generation sequencing (rt-mNGS) for the detection of pathogens in respiratory samples.

**Method:** We used rt-mNGS, using the Seqstant LiveGene Analysis platform, on 335 respiratory samples in comparison to conventional culture results.

**Results:** We observed an overall good concordance in 71.64% (240/335) of the methods. The rt-mNGS outperformed the gold standard culture in 16.12% (54/335) of the samples, while the culture was superior in detecting the clinically relevant pathogen in 12.24% (41/335) of the samples. The non-inferiority of rt-mNGS was statistically significant (δ = 10, *α*= 0.05, 1 *− β*= 0.8). We also observed that the real-time analysis of NGS data is beneficial in obtaining reliable timely results as the initial report at cycle 46 exhibits a Positive Predictive Value (PPV) of 93.75% at the species-level with a sensitivity of 32.09%.

**Conclusion:** Overall, our study showed the non-inferiority of rt-mNGS compared to the standard-of-care microbiology for respiratory samples with statistical significance. Moreover, the rt-mNGS method exhibited superior sensitivity and superior overall performance. It also uniquely detected certain organisms that are typically hard to culture. However, rt-mNGS reported a higher number of false positives and faced limitations in detecting *Aspergillus* spp. In conclusion, the study highlights the potential of rt-mNGS as a powerful tool in clinical diagnostics of respiratory infections and beyond.

## Introduction

Respiratory infections exhibit a multifaceted etiology. While hospital-acquired lower respiratory infections are predominantly caused by bacteria, community-acquired cases may result from an array of pathogens, encompassing predominantly bacteria and viruses, but in some cases also fungi and parasites(1). The intricacy of these infections poses challenges to conventional culture-based microbiological diagnostics, as they may not offer a comprehensive understanding of the causative agents. Conventional culture-based methods may be limited in detecting fastidious and non-viable pathogens, and their sensitivity can be compromised by prior antibiotic exposure. To address these constraints, next-generation sequencing (NGS) technology has emerged as a promising new technique that may complement conventional diagnostic methods, especially for infections in complex clinical situations of immunocompromised patients(2, 3).

NGS-based assays may enhance pathogen detection accuracy, enabling healthcare professionals to identify a broad spectrum of pathogens causing respiratory infections(2, 4, 5). NGS enables the simultaneous sequencing of genetic material from multiple samples, facilitating bioinformatic analysis for comparison against a database to identify potential pathogens(6). Targeted NGS, focusing on specific genetic targets, can enhance sensitivity compared to untargeted approaches, although it may overlook unexpected pathogens or those that are not included in the test panel(2). Therefore, metagenomic next-generation sequencing (mNGS), an untargeted method involving sequencing any nucleic acids, might be a more suitable alternative. However, mNGS may demand more resources than targeted NGS, and sensitivity could be impacted by sequencing depth, potentially reducing microbial reads available for analysis(7). Furthermore, timely microbiological diagnostic reports are critical for informed antimicrobial therapy decisions, optimizing outcomes. However, conventional diagnostics may be time-consuming, and while mNGS holds promise, its turnaround time remains a concern(8, 9). Moreover, the need for bioinformatics expertise can hinder the integration of NGS into routine diagnostics(9).

In that context, the Seqstant LiveGene Analysis platform (Seqstant GmbH, Germany) addresses these challenges by being locally installed directly on sequencers, allowing access to sequence data during the sequencing process. LiveGene Analysis conducts an automated, cloud-based real-time analysis of mNGS data, providing preliminary results before the sequencing run’s completion. This real-time approach significantly reduces the turnaround time for mNGS. This study assessed the performance of Seqstant LiveGene Analysis on a real life setting from respiratory material comparing real-time mNGS (rt-mNGS) data analysis on the Illumina sequencing platform to conventional microbiology testing. The study aimed to (i) compare the performance of conventional microbiological diagnostics with rt-mNGS using a non-inferiority study design, (ii) assess the overall performance and in particular the reliability of preliminary rt-mNGS analysis compared to the final output and post-run analysis methods, and (iii) evaluate the benefits and limitations of the rt-mNGS approach with regards to specific pathogens that are considered to be of clinical relevance.

## Methods

### Study Design

We analyzed a total of 384 left-over respiratory tract samples (bronchial lavage, bronchial secretion, sputum or tracheal secretion) from patients seeking medical care in a tertiary care hospital with standard-of-care microbiology testing and the rt-mNGS approach. Out of the initial 384 samples, 49 were excluded, leaving 335 samples for evaluation. The reasons for exclusion varied, including massive underclustering of the complete sequencing run (16 samples), underclustering of individual samples (8 samples), failed sample preparation (20 samples), and deviations in the evaluation of results such as a reasonable suspicion of sample interchange (5 samples). The analysis of the data is based on pairwise comparison making it independent of patient sex or age and hospitalization status.

### Standard-of-care microbiology testing

Respiratory material was processed according to microbiological quality standards. Testing methods included Gram stain, aerobic culture, anaerobic culture and specific PCR for non-culturable pathogens according to the requested analyses. The culture was performed using commercially available culture media. Species identification was performed by MALDI-TOF MS (Bruker GmbH, Germany) using a score of ≥2.0 for reliable identification to the species level.

### Next generation sequencing

DNA extraction with depletion of human host DNA has been performed with the commercial Qiagen QIAamp DNA Microbiome Kit (QIAGEN GmbH) according to the manufacturer’s instructions. Library preparation has been performed with the IDT Lotus DNA Library Prep Kit (Integrated DNA Technologies, Inc.) with 15 PCR cycles and Seqstant LiveGene sequencing adapters to enable live analysis of the sequencing data.

DNA sequencing was performed on three different devices: Illumina MiSeq, Illumina MiniSeq and Illumina NextSeq 500. The sequencing protocol included 1×151bp (single-end), without the need for additional barcode sequences as inline barcodes are included in the Seqstant LiveGene Analysis adapter design. The target read count for each sample was 1.5M-2M reads, allowing a total of 16 samples on Illumina MiSeq device (target capacity of 25M reads; MiSeq Reagent Kit v3 (150-cycles)), 16 samples on MiniSeq devices (target capacity of 25M reads; MiniSeq High Output Reagent Kit (150-cycles)), and 64 samples on the Illumina NextSeq 500 device (target capacity of 130M reads; NextSeq 500/550 Mid Output Kit v2.5 (150 Cycles)).

The fully automated upload of data to the Seqstant LiveGene Analysis platform was performed by the LiveGene Data Uploader, a software that was installed directly on the used sequencing devices. Once the first Illumina base call files were uploaded to the LiveGene Analysis platform, the live analysis of the data started automatically, adapted from the HiLive2 algorithm (10, 11) and included all required preprocessing and data aggregation steps for pathogen identification. The database contained reference sequences of 509 bacterial and fungal species that are expected to be found in human respiratory tract samples. Analysis results were reported for six different time points (after sequencing cycles 46, 56, 66, 91, 116, 151). The technical workflow from sequencing to results with LiveGene Analysis is shown in Figure 1 and detailed in the supplementary material.

**Figure 1.**
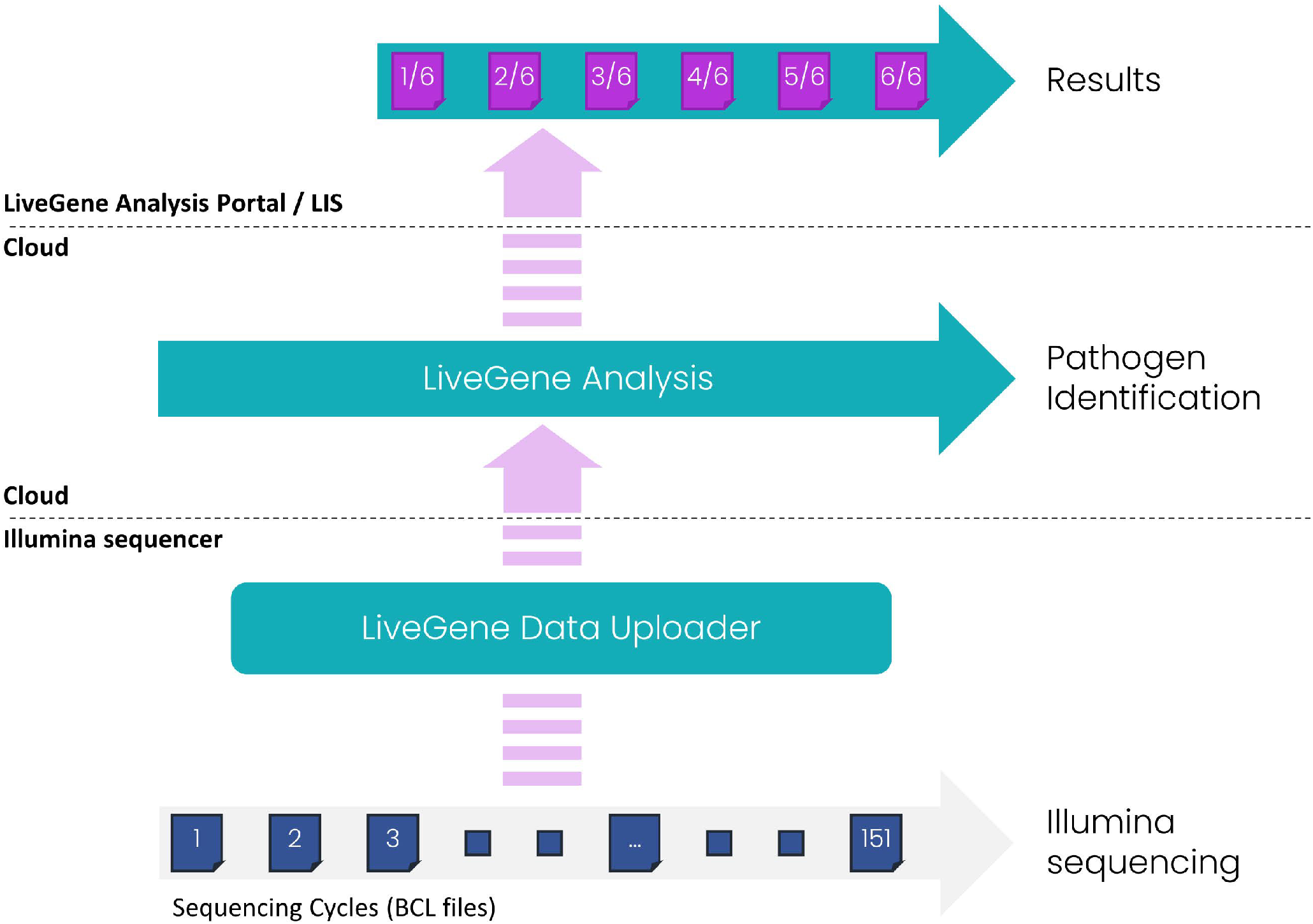
Real-time analysis workflow for rt-mNGS data using the commercial Seqstant LiveGene Analysis software. The LiveGene Data Uploader serves as a conduit for streaming Illumina sequencing data to the cloud platform. This data is subjected to real-time analysis, providing immediate insights even while the sequencing process is ongoing. The LiveGene Analysis Portal allows access to five intermediate reports (generated at cycles 46, 56, 66, 91, 116) and a comprehensive final report. Additionally, the LiveGene Data Uploader can automatically re-integrate the final results, facilitating seamless import of these results into a Laboratory Information System (LIS).

### Cross-validation of hits from LiveGene Analysis

Hits that have been identified only by rt-mNGS were verified as followed: First, the complete preprocessed sequencing data of all samples have been analyzed with Kraken 2(12), using the prebuilt Standard PlusPF-16 database (version: March 14, 2023) and stricter parameters than the default settings and filter parameters (*--confidence* 0.2, at least 50% or 1,000 distinct minimizers, minimum of 5 reads assigned). Additionally, the evaluation of distinct minimizers obtained from the *--report-minimizer-data* option of Kraken 2 corresponds to the HLL-based functionality of KrakenUniq. For species-level classification, species identified with rt-mNGS were only considered confirmed by Kraken 2 if no unidentified other species of the same genus was reported by Kraken 2 with a higher read count than the species identified with rt-mNGS. To account for differences in the database composition and algorithmic characteristics, a second validation step was performed using BLAST(13). Thereby, only the preprocessed reads assigned to an identified organism have been mapped to the nt database with BLAST (downloaded on April 24, 2023, from the NCBI FTP server). The organism was considered confirmed by BLAST if the corresponding taxonomy ID was the ID with the most reads assigned with BLAST. For both tools, Kraken 2 and BLAST, unclassified species were excluded from the evaluation. All identifications with rt-mNGS that were not confirmed by cultivation, Kraken 2 or BLAST were evaluated as false positive hits, regardless of potential differences in the reference databases used or the similarity of differing reported species.

## Results

### Overall comparison rt-mNGS vs culture

A total of 1,139 species occurrences were detected using both rt-mNGS and culture, with 641 considered clinically relevant by expert consensus (Figure 2A). At the species level, 28.97% (270) were concordantly identified by both methods, while rt-mNGS uniquely identified 47.42% (442), and culture identified 23.60% (220). At the genus level, concordance increased slightly to 34.46% (326), with rt-mNGS detecting 43.02% (407) exclusively and culture detecting 22.52% (213). Among 255 clinically relevant species, several species were detected only by rt-mNGS such as *Acinetobacter junii, Fusobacterium nucleatum* and *Tropheryma whipplei*. rt-mNGS was more sensitive than culture to detect pathogens such as *Enterococcus faecium/faecalis, Haemophilus parainfluenzae* and *Streptococcus pneumoniae*. In 335 samples, rt-mNGS identified 89 relevant species missed by culture, indicating culture failed to detect at least one pathogen in over 25% of cases. However, rt-mNGS failed to detect *Aspergillus fumigatus* in five samples and *Aspergillus niger* in one sample and was generally less sensitive regarding yeast and fungi compared to culture (Figure 2B).

**Figure 2.**
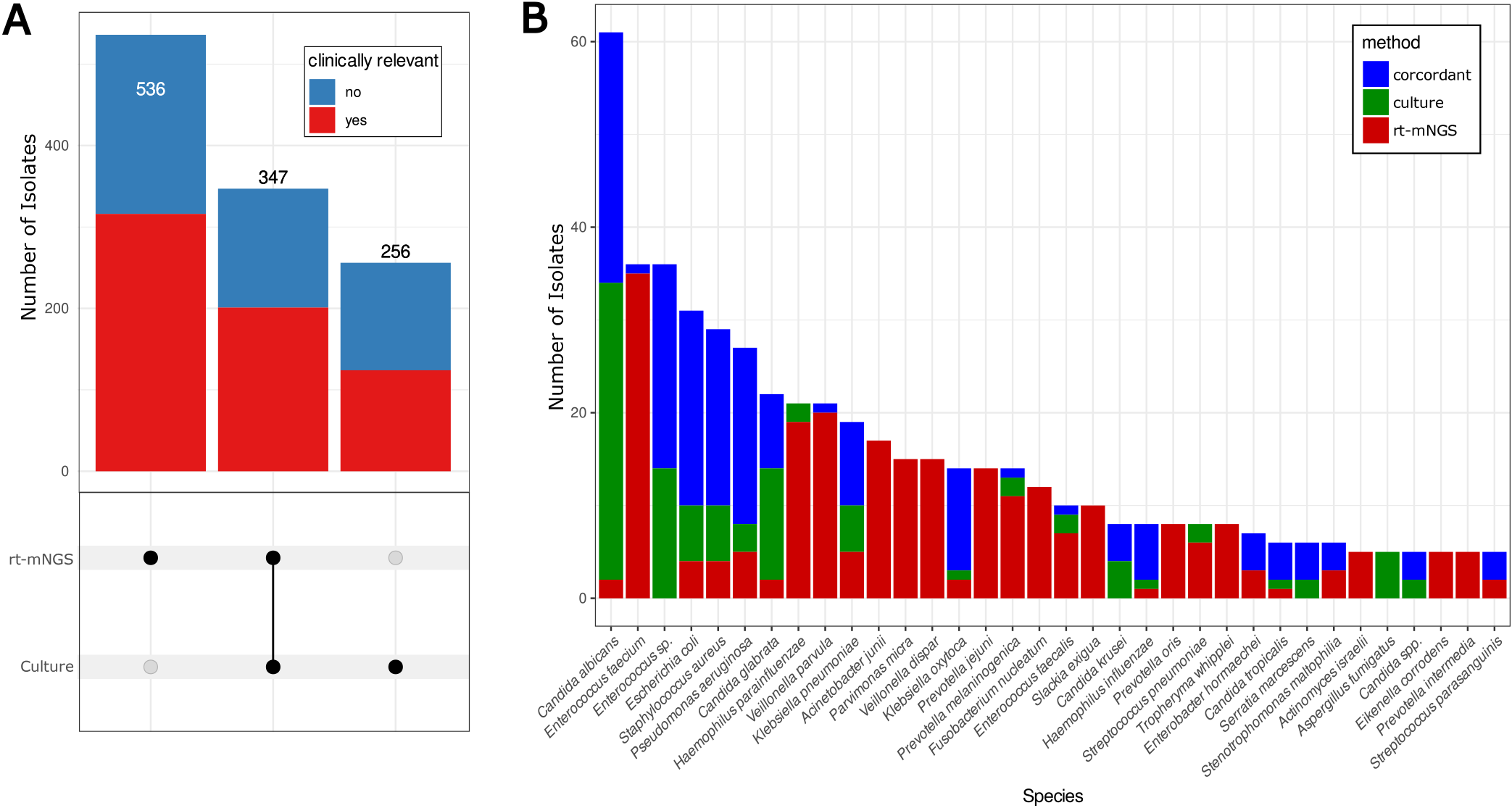
Detection of clinically relevant pathogens by conventional culture and mNGS. **A**. UpSet plots showing the number of species detected in all the sample stratified by clinical relevance (color-coded blue/red) and detection method. **B** Barplot showing the proportion of detection by each method stratified by species. Only clinically relevant species detected more than 5 times in the full study are represented.

Both methods detected relevant pathogens case-wise in 71.64% (240/335) of samples. rt-mNGS outperformed culture in 16.12% (54/335) of cases, while culture was superior in 12.24% (41/335). The non-inferiority of rt-mNGS was statistically significant, with a non-inferiority limit of 10% (δ = 10), significance level of 5% (*α*= 0.05), and statistical power of 80% (1 *− β*= 0.8). rt-mNGS exhibited higher sensitivity at the genus (75.08% vs. 60.18%) and species levels (76.38% vs. 53.61%), particularly for clinically relevant microbes (82.86% vs. 63.67%), though it had a lower Positive Predictive Value (PPV) at genus (96.19% vs. 100%), species (88.64% vs. 100%), and clinical relevance levels (95.31% vs. 100%). This is partly due to the assumption that all positive culture results are true, resulting in a PPV of 100% for culture. The PPV of rt-mNGS improved with higher levels of evidence (LoE), rising from 77.21% at LoE 2 to 93.47% at LoE 4 (Supplementary Table 1). When classifying samples as positive or negative based on the presence of clinically relevant organisms, rt-mNGS matched or exceeded culture across parameters like sensitivity (83.44% vs. 78.53%), Negative Predictive Value (86.43% vs. 83.09%), and overall accuracy (91.94% vs. 89.55%) (Supplementary Table 2).

### Comparison of interim and final reports

We evaluated 817 species occurences (210 clinically relevant) from the 335 patient samples in the comparison of interim and final results provided by Seqstant LiveGene Analysis. The initial report (cycle 46) exhibits a species-level sensitivity of 32.09%, which rises to 62.89% in cycle 91, and peaks at 76.38% in the final report. In regards to clinically relevant species, rt-mNGS exhibits a higher sensitivity starting at 41.39% in the first report (cycle 46), rises to 71.31% in cycle 91, and peaks at 82.86% in the final report, as detailed in Supplementary Table 3.

We also observed that the species-level Positive Predictive Value (PPV) tends to be higher in early reports than in the final report. For instance, the initial report (cycle 46) exhibits a species-level PPV of 93.75%, which decreases to 91.73% for cycle 91 and 88.64% in the final report. The PPV for clinically relevant pathogen starts at 100% for the first and second reports (cycle 46 and 56) and decreases to 98.86% for cycle 91 and 95.31% in the final report (Supplementary Table 3).

We investigated the reliability of interim results at the level of evidence in interim reports and observed a significant loss of evidence in a few cases when multiple species were reported for the same genus due to reporting only the species of the highest evidence for each genus. Therefore, suboptimal species hits may disappear from the final report if the level of evidence for some species increases more than others over time.

### Performance equivalence between devices

The performance of the rt-mNGS method was evaluated across three different sequencing devices: Illumina MiSeq^TM^, MiniSeq^TM^, and NextSeq^TM^, using 64 samples from a single library preparation. These samples contained 42 clinically relevant species occurences. In a pairwise comparison, no clinically relevant pathogens showed a difference in the level of evidence greater than “1” across any devices (acceptable deviation), indicating high consistency. However, 2.4%-3.5% of all 287 identified species showed larger differences. The highest agreement was observed between Illumina MiSeq^TM^ and MiniSeq^TM^ (64.36% agreement; 97.58% acceptable), followed by MiSeq^TM^ and NextSeq^TM^ (65.40% agreement; 96.54% acceptable). The lowest overlap was between MiniSeq^TM^ and NextSeq^TM^ (55.02% agreement; 96.54% acceptable). While read counts had a strong influence on the level of evidence (Rho=0.91, p-value < 0.001), no major differences in absolute read count or case-wise reads were found (Figure 3). The detailed numbers for the individual read counts can be found in the supplementary material.

**Figure 3.**
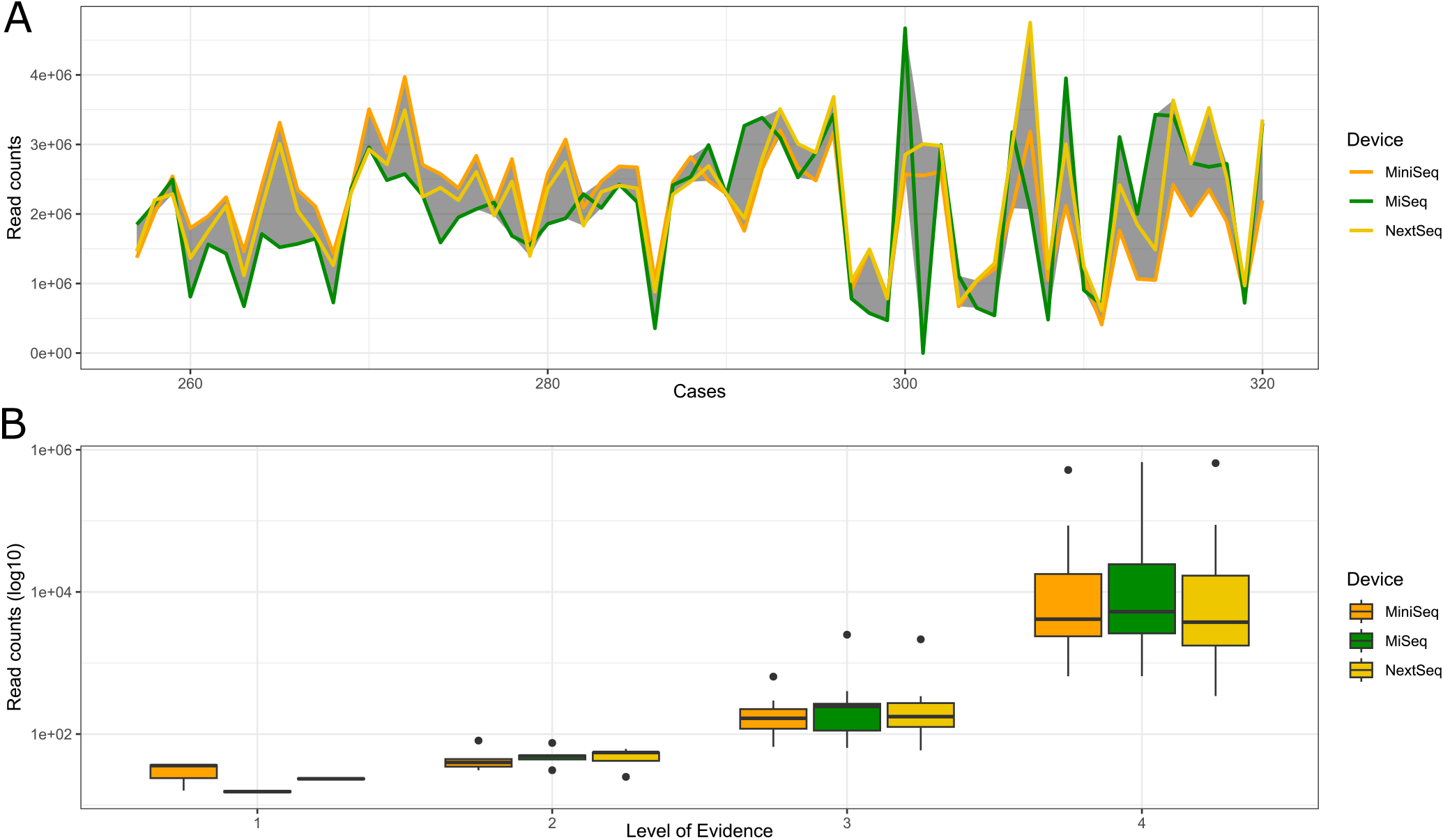
Comparison of the performance in three different Illumina sequencing devices. **A**. The amount of reads per samples for the 64 samples included. The gray area shows the difference between the minimum and maximum read count of the three devices. **B**. Impact of the number of reads counts for each clinically relevant pathogen detected (n=42) on the Level of Evidence.

## Discussion

The current study aimed to compare the performance of rt-mNGS to culture-based microbiological diagnostic in respiratory samples. We were able to demonstrate the non-inferiority of rt-mNGS with statistical significance (δ=10, α=0.05, 1-β=0.8) compared to culture. Generally, rt-mNGS exhibited higher sensitivity but also reported more false positives. This trend aligns with findings from previous studies(5, 14-17). Notably, rt-mNGS uniquely identified certain organisms that are difficult to culture or overlooked in culture media selection. This supports earlier studies that highlighted the superior performance of sequencing-based methods for detecting organisms like *Tropheryma whipplei(17)*. Therefore, the untargeted approach of rt-mNGS offers benefits. Furthermore, the real-time functionality of the Seqstant LiveGene software, which provides interim results during sequencing showed great potential to increase the speed of results. More than half of the relevant microbes were identified after just 56 sequencing cycles, approximately 10 hours before the completion of the sequencing process on an Illumina MiSeq device. Even pathogens that were missed by the cultivation approach could be identified early in the sequencing process with rt-mNGS (46% in cycle 66). This significantly reduces the time-to-diagnosis, which is crucial for critically ill patients(18, 19). Importantly, interim reports did not increase the reporting of false positives compared to the final report, indicating their potential use for interim diagnosis.

The results of the present study demonstrate the benefits of rt-mNGS specifically for different types of respiratory sample material from clinical specimens. However, the general principle of this new diagnostic method is transferable to other sample materials and may be useful in other fields where timely diagnostics is crucial while current methods show limitations in speed and/or sensitivity. These may include wound infections(20, 21), sepsis(22), infective endocarditis(23), and others. It should be noted that adaption of the laboratory workflow and sequencing depth may be required to account for different types of samples. Further, the untargeted characteristics of the rt-mNGS method make it a beneficial addition for all types of infections when considering that one of the main factors for diagnostic delays of infectious diseases is that the appropriate specific test was not ordered(24).

We also showed in a subset of 64 samples that the devices used for sequencing were not influencing the results as the MiSeq, MiniSeq, and NextSeq showed only minor differences, suggesting that rt-mNGS results are generally reproducible across different sequencing devices. However, it’s important to note that the same libraries were used for all three devices, so no conclusions can be drawn about the reproducibility of the overall workflow, including the wet-lab protocols for DNA extraction and library preparation. Such steps are most likely the reason leading to the low identification of “difficult to lyze” microbes observed in our study.

We showed that the rt-mNGS approach is limited in identifying *Aspergillus* spp. While cultivation identified five instances of *Aspergillus fumigatus* and one of *Aspergillus niger*, rt-mNGS did not detect any of these. Previous studies have reported similar limitations in sequencing-based approaches for identifying *Aspergillus* spp., potentially due to the thick polysaccharide cell wall and low fungal load in BALF(25). Possible solutions include enhancing DNA extraction protocols. For example, the introduction of an additional mechanical lysis using zirconium beads may be a promising approach to improve the lysis of gram-positive bacteria and fungi. While comparisons and potential improvements of methods exist(26, 27), the use of commercial kits is beneficial for a routine clinical workflow to increase standardization and reproducibility. At the same time, the availability of such kits being capable of extracting DNA from bacteria and fungi while depleting human DNA is limited. Extraction performance for *Aspergillus* spp. and other fungi should be evaluated for current alternatives to the Qiagen QIAamp DNA Microbiome Kit (Qiagen GmbH) that was used in this study. Besides the evaluation and improvement of DNA extraction methods, algorithmic methods could be tested to increase sensitivity for these and other typical low-abundance species. This could be achieved by establishing species-specific identification thresholds to allow more sensitive detection of microbes that are known to be hard to find with NGS. However, such adaptations need to be extensively validated to prevent a significant increase in false positives for affected species.

We also showed that compared to culture, rt-mNGS had a lower positive predictive value (PPV). This is a known limitation of sequencing-based approaches(5, 14-17). However, the rt-mNGS method used in this study provides a level of evidence (LoE) for each reported organism, which proved to be a reliable measure of results. A LoE threshold of 2 yielded a PPV of 88.57% for clinically relevant species, while an LoE threshold of 4 achieved a PPV of over 99%. This suggests that adjusting the LoE threshold for positivity can rebalance sensitivity and PPV based on the specific case.

## Conclusion

Our study emphasizes the precision of rt-mNGS in identifying pathogens in clinical respiratory specimens, showcasing its higher sensitivity compared to traditional plate cultivation methods. It excels at detecting difficult-to-culture pathogens like *Tropheryma whipplei*, and its interim reporting capability significantly reduces diagnostic turnaround times by 12-24 hours. This makes it especially advantageous for critically ill or immunocompromised patients. Additionally, its high sensitivity positions rt-mNGS as an effective tool for follow-up investigations in culture-negative cases. The study underscores its potential as a transformative diagnostic tool, while calling for further research to improve detection of atypical pathogens and enhance sensitivity for “tough-to-lyze” pathogen such as *Aspergillus spp*..

## Data availability

All the sequencing data are available following the link https://doi.org/10.5281/zenodo.14447316

### Transparency Declaration

The corresponding author affirms that this manuscript is an honest, accurate, and transparent account of the study being reported; that no important aspects of the study have been omitted; and that any discrepancies from the study as planned have been explained.

### Ethics declaration

The local ethics committee approved the study protocol and waived informed consent (S-062/2022).

### Consent for publication

Not applicable.

## Author Contributions

Henri Knobloch, Georg Wrettos, Dennis Nurjadi, Sébastien Boutin, Sabrina Klein, Gerold Untergasser and Tobias P. Loka developed the study design. Georg Wrettos organized the study logistics. Sébastien Boutin, Sabrina Klein and Dennis Nurjadi selected the samples and extracted cultivation results from the clinical routine diagnostic workflow. Sébastien Boutin and Gerold Untergasser designed and optimized the DNA sequencing workflow. Suzan Jakob optimized the laboratory protocols and performed sample and library preparation. Dennis Nurjadi, Sabrina Klein, Elham Khatamzas performed the clinical evaluation of diagnostic results. Tobias P. Loka, Sébastien Boutin and Dennis Nurjadi drafted the manuscript. All authors revised and finalized the manuscript.

## Acknowledgements

We acknowledge Selina Mayer and Nicole Henny for technical assistance with the sequencing devices.

